# A Systems Mechanism for KRAS Mutant Allele Specific Responses to Targeted Therapy

**DOI:** 10.1101/491621

**Authors:** Thomas McFall, Jolene K. Diedrich, Meron Mengistu, Stacy L. Littlechild, Kendra V. Paskvan, Laura Sisk-Hackworth, James J. Moresco, Andrey S. Shaw, Edward C. Stites

## Abstract

A well-established genotype to phenotype relationship in genomic medicine is that activating KRAS mutations indicate resistance to anti-EGFR agents. We used a computational model of Ras signaling to investigate a confusing exception to this relationship whereby colorectal cancers with one specific, constitutively-active, mutant, KRAS G13D, respond to anti-EGFR agents. Our computational simulations of the biochemical processes that regulate Ras suggest EGFR inhibition reduces wild-type Ras activation in KRAS G13D mutant cancer cells more than in other KRAS mutant cancer cells. The model also reveals a non-intuitive, mutant-specific, dependency of wild-type Ras activation on EGFR. This dependency is determined by the interaction strength between a KRAS mutant and tumor suppressor neurofibromin. Our prospective experiments confirm this mechanism that arises from the systems-level regulation of Ras pathway signaling. Overall, our work demonstrates how systems approaches enable mechanism-based inference in genomic medicine.

## Introduction

Cancer treatment decisions are increasingly influenced by which specific genes are mutated within each patient (Chin et al., 2011). This has been referred to as personalized medicine, precision medicine, and genomic medicine. One example of personalized medicine in cancer involves the use of anti-EGFR agents in colorectal cancer patients. Clinical trials have shown that humanized therapeutic antibodies that target EGFR, like cetuximab and panitumumab, provide a survival benefit to colorectal cancer patients (Jonker et al., 2007; Van Cutsem et al., 2007), and these drugs are now approved and utilized for colorectal cancer patients.

Approximately forty percent of patients with colorectal cancer have an acquired *KRAS* mutation within their tumor (Cancer Genome Atlas, 2012). The Ras GTPases (HRAS, NRAS, and KRAS) serve as key nodes in the EGFR signaling network (Figure 1A). The signals that propagate from Ras to its effectors, like the RAF kinases, during the course of EGFR signaling can also be initiated by constitutively active KRAS mutant proteins. These KRAS mutant proteins are not dependent upon EGFR for their activation. Clinical trials have shown that colon cancer patients with a constitutively active KRAS mutants do not benefit from anti-EGFR agents (Amado et al., 2008; Karapetis et al., 2008). This relationship between EGFR inhibitors, KRAS mutations, and colorectal cancer appears consistent with the conventional understanding of EGFR signaling.

**Figure 1.**
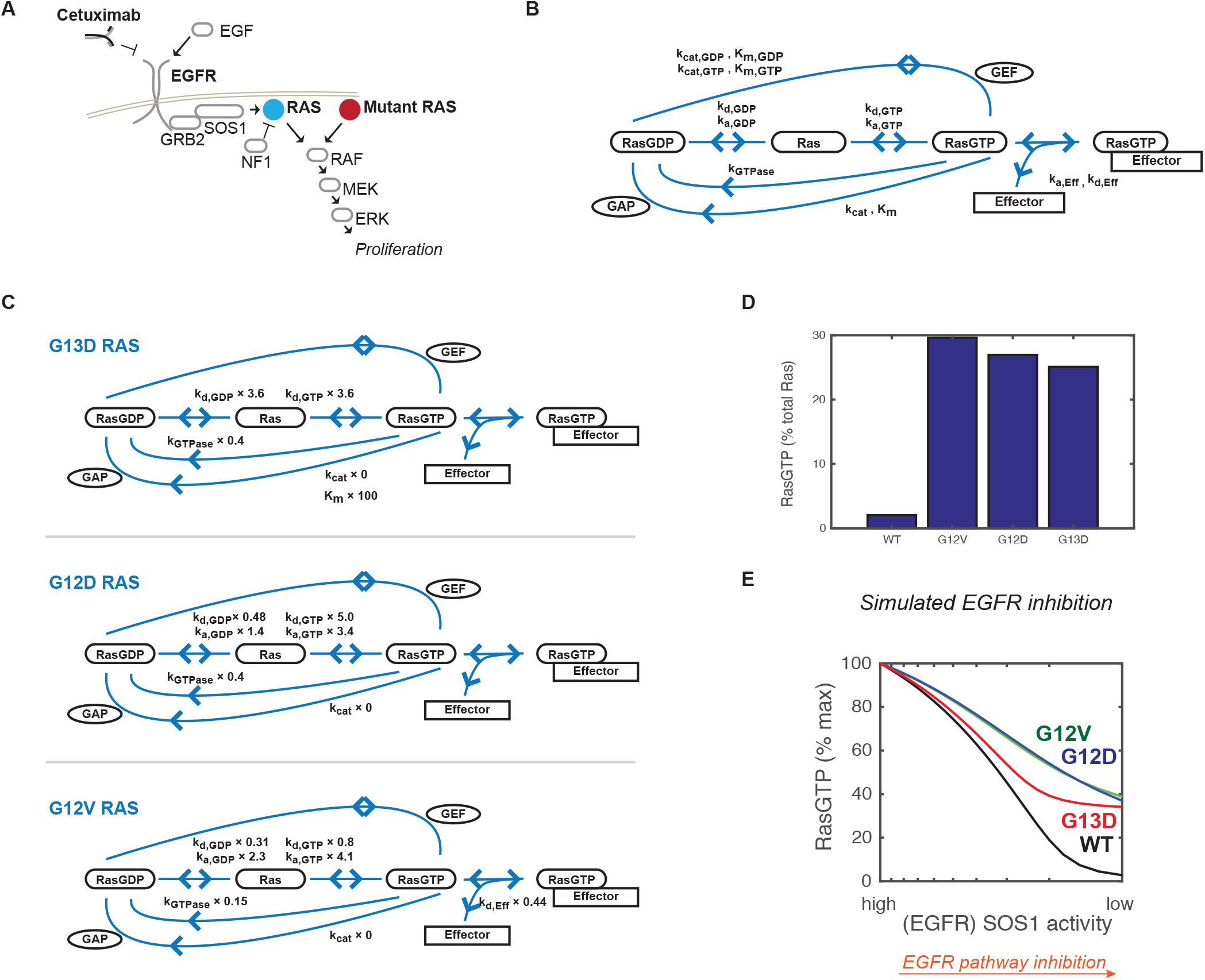
A computational model of Ras signaling emergently reproduces unexplained relationships between oncogenic mutations and response to treatment. **(A)** EGFR signals through the RAS GTPases to drive proliferation. Constitutively active Ras mutants are active in an EGFR-independent manner and are known to cause resistance to EGFR inhibitors. **(B)** The Ras model focuses on the processes that influence the Ras signaling state. The model includes interactions between Ras with GEFs, GAPs, and Effectors as well as slower nucleotide exchange and GTP hydrolysis reactions. For clarity, GTP, GDP, and phosphate ion are not indicated. **(C)** For oncogenic RAS mutants G13D, G12D, and G12V, the network is organized identically, but the specific parameters of the reaction rates can differ from the wild-type value. Parameters that differ between each mutant and wild-type are indicated; the value shown is the factor by which the indicated parameter of the mutant differs from the wild-type value. **(D)** Level of RasGTP within the modeled Ras network for basal conditions with low GEF (SOS1) activity for networks where all Ras is wild-type (WT), or when one modeled allele is Ras G12V, G12D, or G13D. **(E)** Simulated anti-EGFR dose response from the computational Ras model.

Several studies suggest that the relationship between oncogenic KRAS mutants and the response to EGFR inhibitors is more complicated. Initially, a retrospective analysis of phase III clinical trial data found that the anti-EGFR agent cetuximab benefited colorectal cancer patients with a *KRAS G13D* mutation, but not patients with any other *KRAS* mutation (Figure 1B) (De Roock et al., 2010). Although this claim has been further supported with additional clinical trials and experimental model systems (De Roock et al., 2010; Nakamura et al., 2017; Tejpar et al., 2012), the finding remains controversial as expert opinion has been that it is difficult to reconcile known principles of Ras biology with *KRAS G13D* patients responding differently (Morelli and Kopetz, 2012; Peeters et al., 2013; Segelov et al., 2016; Stephen et al., 2014). Without a mechanism, expert opinion has been to assume that the *KRAS G13D* mutation confers resistance and to consider it a contraindication to anti-EGFR agents just like the other constitutively active *KRAS* mutants. Resolving this problem has the potential to benefit a large number of cancer patients. For example, there are approximately ten-thousand new cases of *KRAS G13D* colorectal cancer in the United States alone (Gao et al., 2013; Siegel et al., 2018).

Here, we describe our computational and experimental investigation of this problem. We first applied our computational systems biology methods for studying Ras mutant proteins to determine whether these methods could provide any new insights into this problem. We found that the controversial KRAS G13D behavior that has been interpreted to be inconsistent with known mechanisms of Ras biology is actually fully consistent with known mechanisms of Ras biology. Our model suggests that cancers with the G13D mutant are more sensitive to EGFR inhibition because levels of active, cellular, wild-type RasGTP decrease in G13D cancers much more than in cancers with other Ras mutations. Our experiments confirm this predicted difference. Our model also suggests that the key difference between G13D and the other common Ras mutants is that G13D does not bind well to the tumor suppressor neurofibromin. Our experiments confirm this mechanism. Overall, this work demonstrates the power of computational systems biology approaches for problems in personalized medicine, and it also highlights the necessity of computational methods as a tool for understanding the behaviors of biological networks important to disease.

## Systems Modeling of Oncogenic KRAS mutants

We previously developed a mathematical model of the processes that regulate Ras signaling based upon the well-established architecture of the Ras signaling module and the available biochemical rate constants of wild-type and mutant proteins (Stites et al., 2007) (Figure 1B). A Ras mutant is incorporated into the computational model through the inclusion of its specific biochemical rate constants. We use model simulations to find the behaviors that logically follow from this well-accepted information, but may nevertheless be non-obvious due to the complexity and scope of the system (Stites and Shaw, 2018; Stites et al., 2015; Stites et al., 2007).

Here, we utilize our mathematical model to computationally investigate how Ras mutations should influence the response to EGFR. The three most common Ras mutants in colorectal cancer are G12D, G12V, and G13D (Cancer Genome Atlas, 2012). We updated our model, which already included G12D and G12V mutants (Stites et al., 2007), to also include the G13D mutant by incorporating the known biochemical differences between each mutant and wild-type Ras, as has been previously measured experimentally (Gremer et al., 2008; Palmioli et al., 2009) (Figure 1C). We found that the available data for the G13D mutant were sufficient to result in the constitutive activation once they were applied to the model, just as the available data for G12D and G12V have been shown to be sufficient to explain these mutants’ constitutive activation (Figure 1D).

We then used the model to investigate how systems with each mutant would respond to EGFR inhibition. We did this by using the computational model to find the levels of total, cellular, active RasGTP that should occur for conditions of high EGFR activation (which leads to Ras activation through the RasGEF Sos1) to conditions of low EGFR activation (where low levels of Ras activation by the RasGEFSos1 would occur). Surprisingly, our simulations of EGFR inhibition, which were based on biochemical data of these mutants, found the G13D-containing network displayed larger reductions in Ras signals than the G12D and G12V-containing networks (Figure 1E). This was notable, because expert opinion had been that it did not make sense for different Ras mutants to respond differently to EGFR inhibition. Our analysis revealed that it is fully consistent with known mechanisms of Ras signaling for some mutants to respond more strongly to EGFR inhibition. Moreover, our analysis suggests that the available biochemical data are sufficient to explain a mechanism by which G13D would be the most sensitive of the most common KRAS mutants in colorectal cancer. This highlights the limitation of expert level intuition to logically deduce the behaviors of complex nonlinear networks, like the Ras network.

## Evaluation of an Experimental Model System for this Phenomenon

To experimentally study KRAS-allele specific differences and specific predictions of the model, we obtained a panel of isogenic colorectal cancer cells (De Roock et al., 2010) that was previously derived from the SW48 colon cancer cell line and used to study the *KRAS G13D* response to cetuximab. We obtained isogenic cells with the following *KRAS* genotypes: *G12D/WT* (*G12D* cells), *G12V/WT* (*G12V* cells), *G13D/WT* (*G13D* cells), and *WT/WT* (*WT* cells) (Figure 2A). The mutant isogenic cells display constitutively elevated levels of active RasGTP when compared to the parental *WT* cells (Figure 2B), consistent with all three of these mutants being constitutively active.

**Figure 2.**
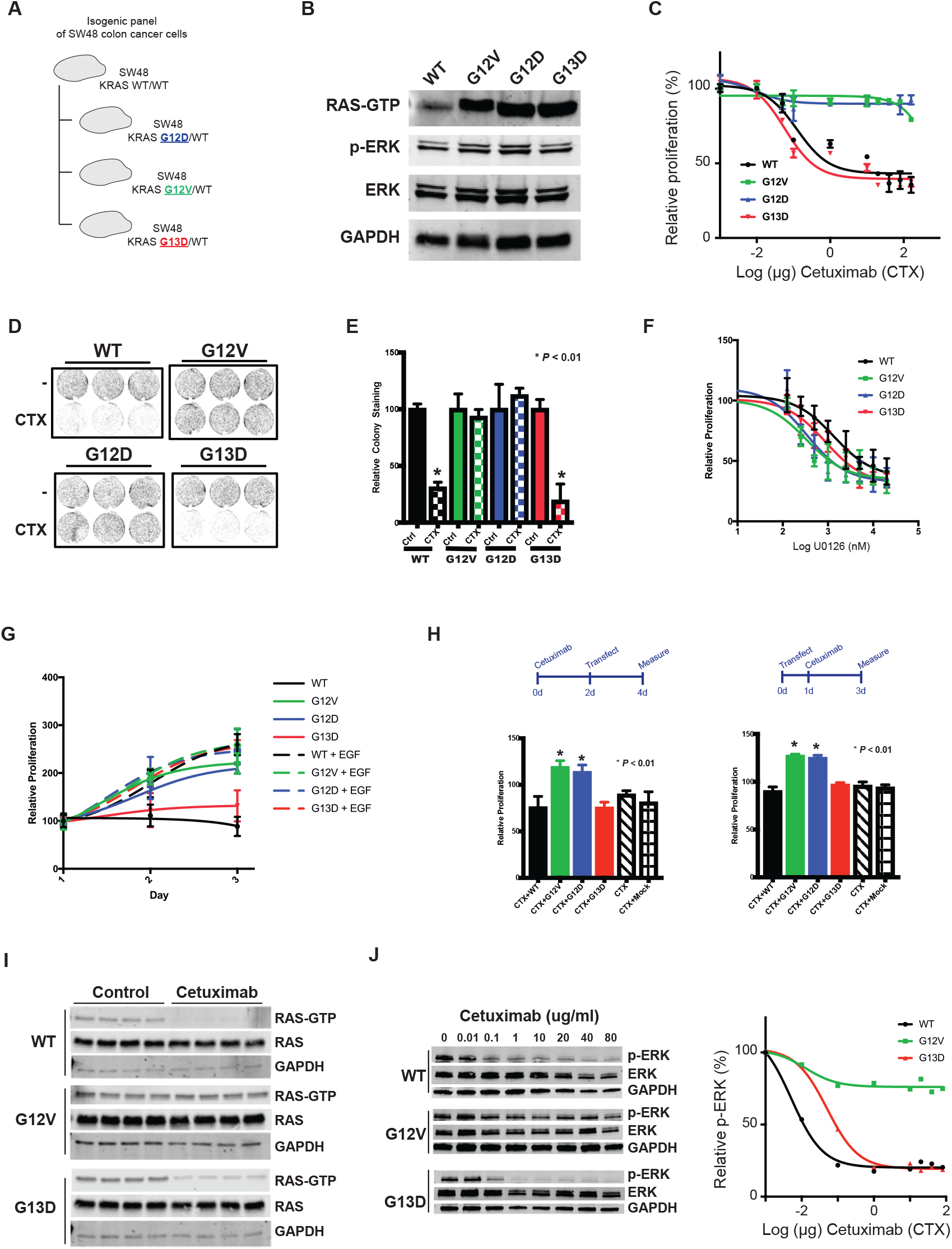
An experimental system that recapitulates the KRAS mutant-specific response to anti-EGFR agent cetuximab in colorectal cancer. **(A)** A panel of isogenic cells derived from *KRAS WT* SW48 colon cancer cells to express one each of the three most common KRAS mutations in colon cancer (*KRAS G12D, KRAS G12V*, and *KRAS G13D*). **(B)** Immunoblots of isogenic cell lysates, including RasGTP pulled down with a Ras Binding Domain (RBD), demonstrate the constitutive activation of these mutants. **(C)** Cetuximab (anti-EGFR) drug dose responses for *KRAS WT* SW48 (*WT*) colon cancer cells and three derivative isogenic cell lines. **(D)** Colony formation assay for each cell line in the isogenic panel. **(E)** Quantification of the colony formation assay for each cell line in the isogenic panel. **(F)** U0126 drug dose response for each cell line in the isogenic panel. **(G)** Proliferation time course for the isogenic panel grown in low serum media and in low serum media with supplemental EGF. **(H)** Proliferation of KRAS WT SW48 cells transfected with KRAS G12V, G12D, G13D and treated with cetuximab. (left) Cells were treated with cetuximab before transfection. (right) Cells were transfected before treatment with cetuximab. **(I)** Ras binding domain (RBD) pull-down Ras activation assays for isogenic SW48 cells grown without and with cetuximab. Four technical replicates for each condition were included in each experiment. **(J)** Immunoblots of ERK phosphorylation for isogenic SW48 cells grown in different concentrations of cetuximab (left); quantification of relative phospo-ERK levels (right).

We performed dose response experiments with the anti-EGFR agent cetuximab to evaluate described difference for these cells. When treated with increasing doses of the anti-EGFR agent cetuximab, both the *G13D* cells and *WT* cells displayed reduced proliferation (Figure 2C) and reduced colony formation (Figure 2D, E) whereas the *G12D* and *G12V* cells were unaffected. We also evaluated dose responses to MEK inhibitors to evaluate whether these cells were more sensitive to any inhibition of the pathway. We observed that all cell lines responded similarly to MEK inhibition (Figure 2F), suggesting that the *G13D* cells are not simply more sensitive to all agents that target the EGFR/RAS/ERK pathway.

We hypothesized that if there was a difference in how these cells depended upon EGFR signals, that we should be able to detect net proliferation differences when these cells are grown in low amounts of serum. We grew these cells in low serum media, and also in low serum media supplemented with additional EGF. Consistent with our hypothesis, we observed that *G13D* and *WT* cells proliferated more slowly than *G12D* and *G12V* cells when grown in low serum media, but that all cells proliferated at a similar rate when supplemental EGF was added to the media (Figure 2G). This further suggests *G13D* cells display an increased dependency upon EGFR signaling compared to *G12D* and *G12V* cells. As the patterns of response appears analogous to the clinical observations regarding *KRAS* genotype and response (De Roock et al., 2010), it suggests this cell line would be useful to test our experimental model.

## An alternative experimental model of resistance suggests G13D is more sensitive to EGFR inhibition

We hypothesized that the introduction of mutant KRAS G12D and G12V into the WT Ras cells should reduce sensitivity to cetuximab, while introduction of KRAS G13D would have a minimal effect on sensitivity. In our experiments, we observed that transfected *KRAS G12D* and *G12V*, but not *G13D* and *WT*, promoted resistance to cetuximab, consistent with our hypothesis and consistent with the G13D mutant being comparably more sensitive to EGFR inhibition (Figure 2H).

## Experimental Evaluation of Predicted Signaling Differences

Our model suggests that there should be signaling differences between *G13D* cells and cells with one of the other common *KRAS* mutations (*G12D* cells and *G12V* cells). We measured levels of active, GTP-bound, Ras (RasGTP) for cells treated and not treated with cetuximab, and we detected a reduction in RasGTP only in *G13D* and *WT* cells, but not in *G12V* cells (Figure 2I). As RasGTP signals are transmitted downstream through the ERK MAPK cascade (Figure 1A), we also measured active, phosphorylated, ERK for cells treated with different doses of cetuximab. We detected reductions in the active, phosphorylated, form of ERK for both the sensitive *G13D* and *WT* cells that had been treated with cetuximab, but not for the resistant *G12V* cells (Figure 2J).

## Model Prediction: Differences are Mediated by Wild-Type Ras

Our computational model includes both mutant (KRAS) and wild-type (KRAS, NRAS, and HRAS) pools of Ras because colorectal cancer cells express all three Ras proteins (Mageean et al., 2015). (We use “*WT*” to indicate the genotype of isogenic cells that have no mutant Ras, and we use “wild-type” to indicate the non-mutant Ras protein that can be found in both *WT* and mutant cell lines.) The differences in total RasGTP that our model predicts are accordingly distributed between GTP bound mutant Ras proteins and GTP bound wild-type Ras proteins.

We queried our model to determine whether the predicted changes in signal were coming from mutant Ras, wild-type Ras, or both. Our simulations suggest that EGFR inhibition should cause no appreciable changes in the level of mutant Ras bound to GTP (Figure 3A). This is consistent with the conventional wisdom that anti-EGFR agents should not influence mutant Ras signaling. However, our simulations predicted that EGFR inhibition should result in large changes in wild-type RasGTP (Figure 3A). This suggests that the non-obvious response to anti-EGFR agents may have a basis in wild-type Ras signaling.

**Figure 3.**
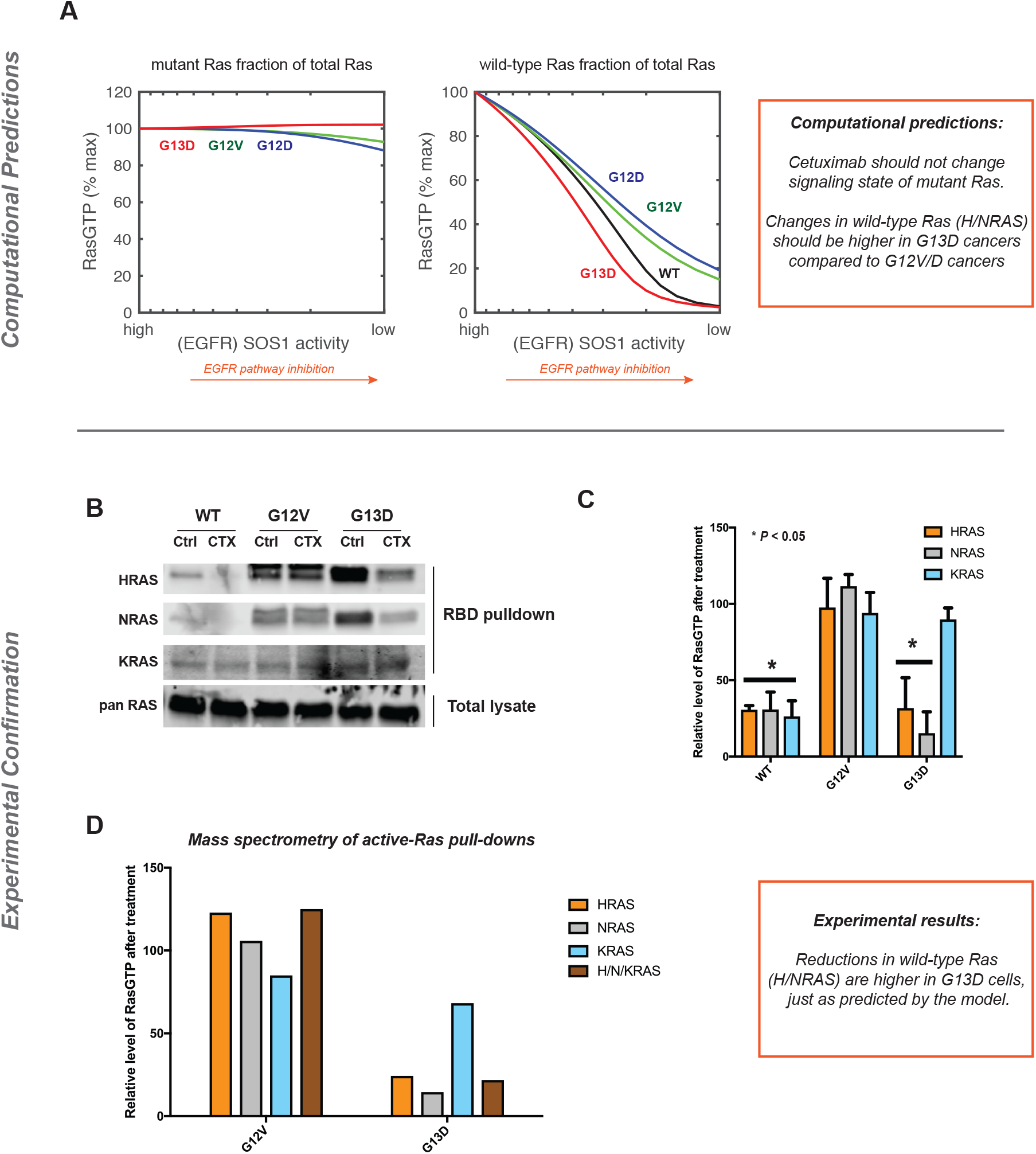
The Ras model predicts, and experiments confirm, that wild-type Ras activation distinguishes sensitive from non-sensitive cancer cells. **(A)** Simulated anti-EGFR dose response for the Ras model, further subdivided to reveal the change in active, GTP-bound Ras mutant (left) and in active, GTP-bound, wild-type Ras (right) within each modeled genotype. **(B)** Ras binding domain (RBD) pull-down Ras activation assays for isogenic SW48 cells grown without and with cetuximab. **(C)** Quantification of the ratio of RasGTP between cetuximab treated and untreated cells for three RBD assays. **(D)** Mass spectrometry quantification of the ratio of RasGTP levels between cetuximab treated and untreated cells for wild-type HRAS, wild-type NRAS, total KRAS, and wild-type H/N/KRAS.

## Experimental Confirmation: Differences are Mediated by Wild-Type Ras

We returned to our experimental system to test the model-based hypothesis that EGFR inhibition causes a larger drop in wild-type RasGTP in G13D cells than in cells with one of the other common Ras mutants. We measured Ras activation in the presence and absence of cetuximab for each of the Ras proteins (HRAS, NRAS, and KRAS) (Figure 3B,C) by using antibodies specific for each form of Ras. We observed a large reduction in wild-type HRAS-GTP and wild-type NRAS-GTP after cetuximab treatment only in *G13D* and *WT* cells, consistent with our model’s predictions. We also observed a larger reduction in KRAS-GTP in *WT* cells than in *G12V* and *G13D* cells, consistent with the presence of one constitutively active *KRAS* allele for the two mutant cell lines.

To complement these studies, we developed a mass-spectrometry assay that could quantify levels of active HRAS, NRAS, KRAS, and H/N/KRAS through the use of isotopically labeled peptides unique to HRAS, NRAS, KRAS, wild-type H/N/KRAS, and the G12V and G13D Ras mutants. This approach revealed greater reductions in active wild-type HRAS, wild-type NRAS, and wild-type H/N/KRAS in G13D cells treated with cetuximab than in G12V cells treated with cetuximab (Figure 3D). Additionally, KRAS in G13D cells displayed a partial reduction, consistent with one KRAS allele being wild-type and one KRAS allele being mutant.

## Computational Hybrid Mutants Show the GAP K_m_ is the Necessary and Sufficient G13D Parameter

The different responses of the Ras mutants in our computational model must follow from the differences in their specific parameters. We therefore computationally dissected the modeled Ras mutants to determine which parameters determine sensitivity to anti-EGFR agents. We did this by creating new computational Ras mutant through the process of mixing the parameters from the G13D, G12V, and G12D mutants, effectively creating computational hybrid Ras mutants (Figure 4A). We used our model to simulate dose responses to anti-EGFR agents for each of 648 different computational hybrids, and then we determined whether any single parameter could distinguish between the sensitive and resistant hybrid mutant networks. Our analysis found that all hybrids that were sensitive to simulated EGFR inhibition contained the K_m_ characterizing the interaction between KRAS G13D and the Ras GTPase Activating Protein (GAP) neurofibromin (NF1), and also that all mutants that were insensitive to simulated EGFR inhibition had the K_m_ value that applied to the G12D and G12V mutants (Figure 4B). Thus, this demonstrates that that this parameter is necessary and sufficient for sensitivity to EGFR inhibition in our systems model of Ras signaling.

**Figure 4.**
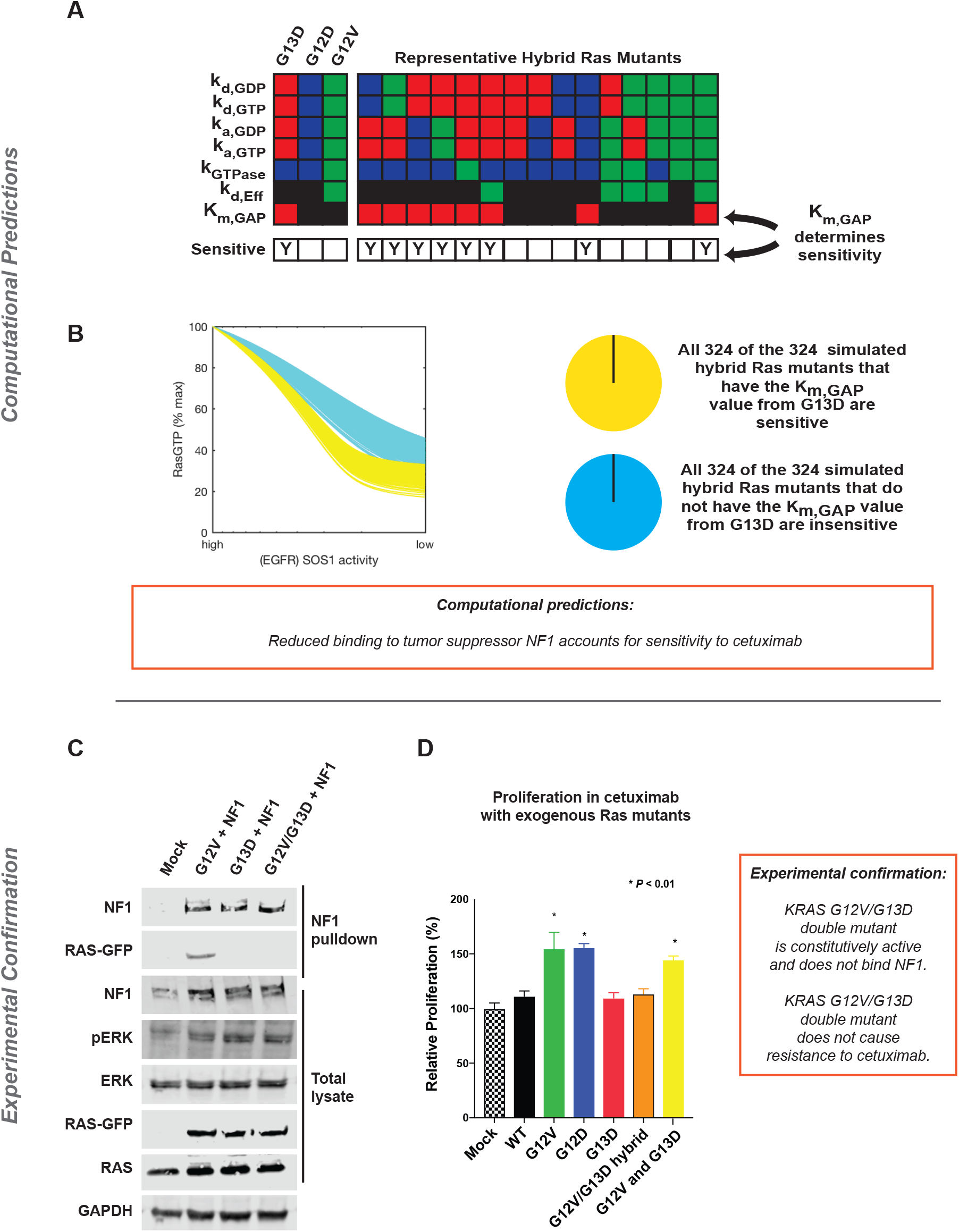
The Ras model predicts, and experiments confirm, that mutant-specific interactions with tumor suppressor NF1 determine whether or not cells respond to anti-EGFR agents. **(A)** Schematic to explain computational Ras hybrid mutants. G13D, G12D, and G12V have been described to differ in seven different biochemical parameters. 648 different computational hybrids were generated by considering all of the possible combinations of these parameters. For each mutant, the model was evaluated to determine whether the computational hybrid was sensitive or resistant to simulated EGFR inhibition. **(B)** Simulated dose responses for all 648 hybrids color coded based on whether the hybrid had the Ras/NF1 K_m_ value of the G13D mutant or of the G12V and G12D mutant. **(C)** Co-immunoprecipitation of NF1 with KRAS G12V, G13D, and G12V/G13D hybrid from mixed, transfected cell lysates. **(D)** Proliferation of cetuximab treated *KRAS WT* SW48 cells transfected with KRAS WT, G12V, G12D, G13D, G12V/G13D hybrid, and both G12V & G12D.

## Experimental Hybrid Mutant Confirms the Importance of the GAP/Ras K_m_

We set out to test the computational results that suggest the strength of the interaction with NF1 can determine whether a mutant is sensitive or resistant to cetuximab. We first created a new hybrid Ras mutant, KRAS G12V/G13D, where the glycine residues at codons 12 and 13 were replaced with a valine and aspartic acid, respectively. We found this mutant was constitutively active, as demonstrated by its ability to lead to increased ERK phosphorylation (Figure 4C). We also found that this combination mutant bound much less well to NF1 (Figure 4C).

If the ability to bind NF1 is the critical factor that determines whether or not a mutant promotes resistance to cetuximab, as suggested by our model, we reasoned that the KRAS G12V/G13D mutant would not promote resistance to cetuximab. We used our transfection-based assay to evaluate the ability of transfected Ras mutants to alter WT cells sensitivity to cetuximab. Consistent with our hypothesis, we observed that the G12V/G12D double mutant did not promote resistance, despite being constitutively active (Figure 4D).

## A Mechanism for KRAS Mutant Allele-Specific Responses to EGFR Inhibition

We considered how differences in the interaction between KRAS and NF1 might result in differences in network signal output. Our previous systems analysis of oncogenic Ras found that the reversible interaction between a Ras mutant and a Ras GAP can promote wild-type Ras activation (Stites et al., 2007), as the GAP-insensitive Ras mutant can effectively behave as a competitive inhibitor of Ras GAPs (Bollag and McCormick, 1991). Our previous prediction that mutant Ras leads to wild-type Ras activation has been reproduced in several other studies (Grabocka et al., 2014; Jeng et al., 2012; Keller et al., 2007; Lim et al., 2008). Our new study suggests that G13D is an exception to this process because it binds much less well to NF1 (Gremer et al., 2008) and therefore cannot lead to wild-type Ras activation via the competitive inhibition of NF1 Ras GAP activity.

We therefore propose a mechanism that explains why KRAS G13D, but not other common KRAS mutants like G12D and G12V, responds to cetuximab (Figure 5). In a *WT* cell, Ras activation is dependent upon EGFR and can be counteracted with EGFR inhibitors. In a *G12D* or *G12V* cell, the mutant KRAS is constitutively active. Through the competitive inhibition of Ras GAP NF1, wild-type Ras is also active in an EGFR independent manner and the cells will be insensitive to EGFR inhibition. In a *G13D* cell, the mutant KRAS is constitutively active and wild-type Ras activation is dependent on EGFR because the G13D mutant cannot drive wild-type Ras activation through the competitive inhibition of Ras GAP NF1. Assuming that the activation of proliferative signals downstream from Ras requires a quantity of Ras signal that is greater than the mutant alone can typically provide, inhibition of wild-type Ras through EGFR inhibition should negatively impact proliferation signals within the *G13D* cell. This assumption that wild-type Ras signaling is required in addition to mutant Ras signaling is consistent with emerging data that cancer promotion requires both wild-type and mutant Ras signals (Grabocka et al., 2014; Jeng et al., 2012; Young et al., 2013).

## Experimental Testing of the Proposed Mechanism

We desired to test and confirm this proposed mechanism. We hypothesized that reduced expression of NF1 would make both *G13D* cells and *WT* cells less sensitive to cetuximab but would not largely affect *G12V* cells. This is because we reasoned reduced NF1 should result in increased wild-type RasGTP, thereby making these cells less dependent upon EGFR for wild-type Ras activation. We performed siRNA knockdown experiments of *NF1* in *WT, G13D*, and *G12V* cells and compared proliferation in the presence and absence of cetuximab. As hypothesized, *NF1* knockdown reduced the sensitivity of *G13D* cells and *WT* cells to cetuximab with minimal effect on *G12V* cells (Figure 6A). We also hypothesized that increased expression of NF1 should make *G12V* cells more sensitive to cetuximab. This is because we reasoned that these cells would become more dependent upon EGFR for wild-type Ras activation as NF1 levels increased. To test, we transfected *WT, G13D*, and *G12V* cells with *NF1* and then treated with cetuximab. As hypothesized, increased *NF1* expression made the *G12V* cells significantly more sensitive to cetuximab (Figure 6B). Lastly, we reasoned that this mechanism also suggests that the introduction of KRAS G13D into a *G12V* or *G12D* cell would not cause the *G12V* or *G12D* cells to become sensitive to cetuximab, as the codon 12 KRAS mutant can still competitively inhibit NF1. We experimentally tested this hypothesis by transfecting *G12V* and *G12D* cells with *KRAS G13D* and found that the introduction of the *KRAS G13D* mutant did not cause the cells to become sensitive to cetuximab (Figure 6C), consistent with our proposed mechanism.

**Figure 5.**
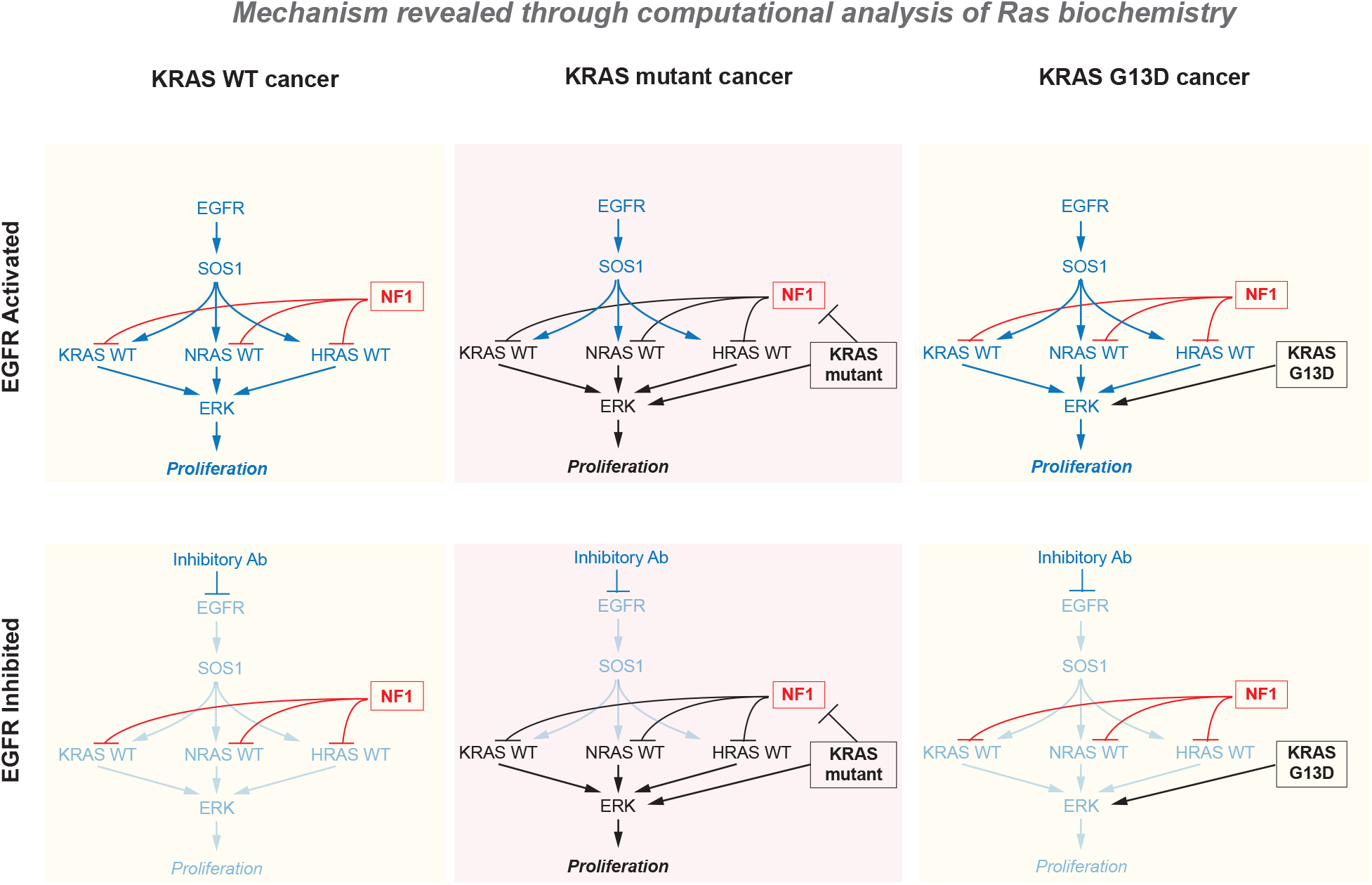
A mechanism for KRAS-allele specific response to anti-EGFR agents. In a *KRAS WT* cancer, NF1 ensures there are low levels of RasGTP when EGFR is not active (or is inhibited). In *KRAS G12D* and *KRAS G12V* cancers, mutant Ras is active. Wild-type Ras is also active through the competitive inhibition of NF1 through the non-productive interaction between these Ras mutants and NF1. In a *KRAS G13D* cancer, mutant Ras is active but wild-type Ras remains dependent on EGFR for activation due to the inability of KRAS G13D to bind NF1.

**Figure 6.**
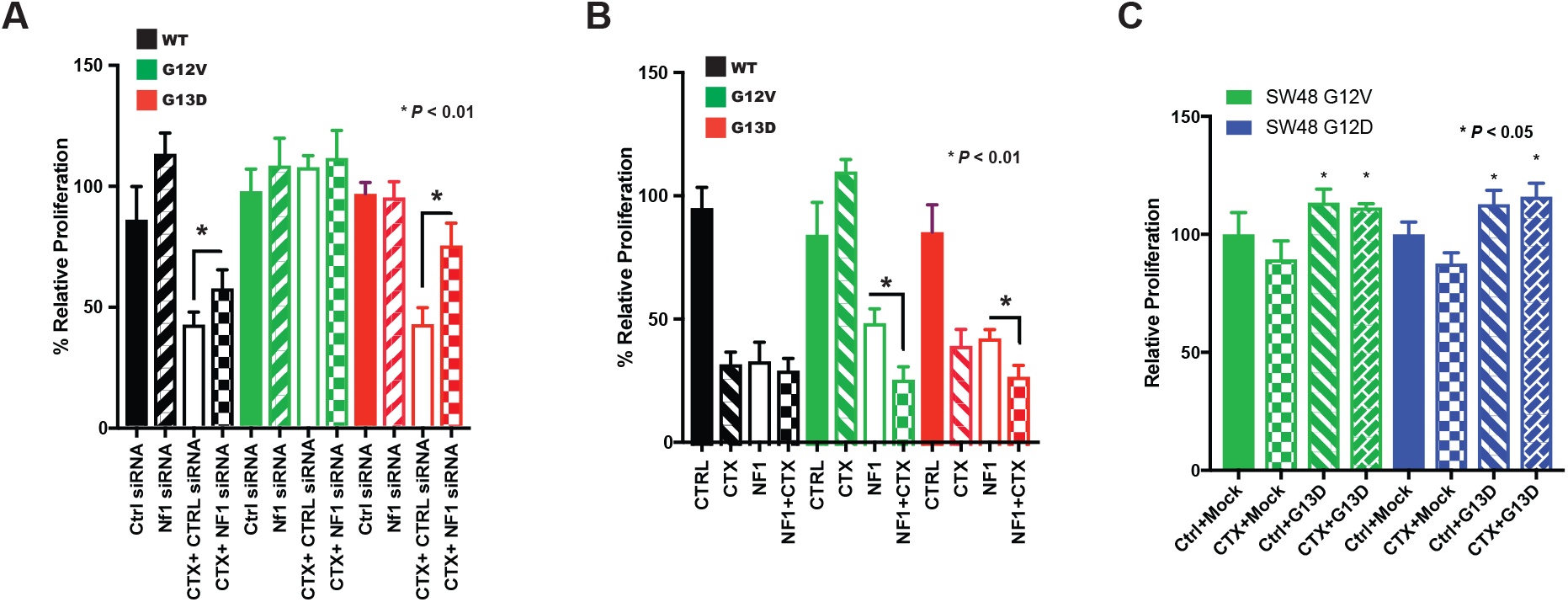
Experimental confirmation of the systems mechanism for KRAS mutant allele specific responses to targeted therapy. **(A)** Proliferation assays for isogenic SW48 cells with siRNA knock down of NF1 and/or with cetuximab treatment. **(B)** Proliferation assays for isogenic SW48 cells with neurofibromin transfection and/or with cetuximab treatment. **(C)** Proliferation of cetuximab treated *KRAS G12V* SW48 cells (left) and *KRAS G12D* SW48 cells (right) transfected with *KRAS G13D.*

## Discussion

Our work demonstrates how systems approaches can uncover non-obvious, mechanistic bases for clinical observations that otherwise defy expert-level explanation. Many genes associated with cancer and other diseases have multiple pathological variants. Our work demonstrates that apparently similar variants can have different responses to the same pharmacological treatment. As clinical genomics becomes more common, and as the number of targeted therapies approved and in development continues to grow, we believe that it will be increasingly necessary to perform integrated mathematical analysis of biomolecular systems to understand how mutant allele-specific behaviors emerge and influence response to treatment.

## Methods

### Mathematical Model and Analysis

Details of the model and its development have been published previously (Stites, 2010; Stites and Ravichandran, 2012a, b; Stites et al., 2007). The model focuses on Ras and the types of proteins that directly interact with Ras to regulate Ras GTP levels: Ras GEFs (e.g. SOS1), Ras GAPs (e.g. neurofibromin/NF1), and Ras effector proteins (e.g. the RAF kinases). The model includes 1) GEF mediated nucleotide exchange, 2) intrinsic nucleotide exchange, 3) GAP mediated nucleotide hydrolysis, 4) intrinsic nucleotide hydrolysis, and 5) effector binding. Reaction mechanisms modeled are presented below and numbered in correspondence with the above list of processes:

1. RasGDP + GEF <-> RasGTP + GEF
2. RasGXP <-> Ras ()
3. RasGTP + GAP -> RasGDP + GAP
4. RasGTP -> RasGDP
5. RasGTP + Effector <-> RasGTP-effector

GXP could indicate GTP or GDP for 2, above. Free GTP, GDP, and P_i_ are not indicated in the biochemical reactions above for simplicity and are approximated to be constants. GEF and GAP reactions, 1 and 3 above, are described mathematically with reversible and irreversible Michaelis-Menten kinetics, respectively. The other reactions are described with first- and/or second-order mass-action kinetics. It is assumed that wild-type and Ras mutant proteins have identical reaction mechanisms as indicated above, and that differences in rate constants (or enzymatic parameters) for the reactions account for described differences. For example, Ras mutant protein G12V hydrolyzes GTP more slowly than does wild-type Ras. In this case, the rate constant for this reaction k_GTPase,G12V_ is smaller than the rate constant for the same reaction with wild-type Ras, k_GTPase,WT_. All reactions are grouped into a set of differential equations and the steady-state quantity of RasGTP-effector complexes (and RasGTP) is solved for the specified conditions.

Parameters of the model for proteins correspond to biochemically observable properties. Approximate concentrations for the total amount of Ras GTPase, effectors, and basally active GEF and GAP have previously been estimated, utilized, and published(Stites and Ravichandran, 2012b). Rate constants and enzymatic properties (e.g. K_m_) for wild-type Ras proteins have been previously obtained, utilized, and published. Mutant proteins can be characterized by their difference from wild-type proteins in terms of a multiplicative factor, α. Values for α are determined from previous experimental studies that measured the desired property for both wild-type and mutant Ras proteins (Chuang et al., 1994; Eccleston et al., 1991). For G12V and G12D, we use the same α values that were previously obtained and utilized in our model (Stites et al., 2007). For G13D previous experiments described this mutant to have an elevated nucleotide dissociation rate compared to wild-type Ras (α = 3.6625) (Palmioli et al., 2009). Previous studies have also described Ras G13D to be insensitivity to Ras GAP (Fischbach and Settleman, 2003), and to have no appreciable binding to the Ras GAP NF1 (Gremer et al., 2008). A 100-fold increase in the K_m_ of GAP on Ras G13D is used to model the immeasurable binding to the Ras GAP NF1. We estimated the change must be at least 100 times large as changes of approximately 50-fold have previously been measured for other Ras mutants (Donovan et al., 2002), so we assumed that the difference must be larger to be undetectable. The decreased GTPase activity of the G12D mutant is used for the G13D mutant because we could not find an α factor at the time we began our; using the same value as G12D allowed us to introduce impaired GTPase activity while also allowing us to focus on the known biochemical differences.

Computational “hybrid” mutants are modeled mutants that have properties of two distinct Ras mutants. For example, a hybrid Ras mutant may be modeled with all of the properties of Ras G12D, except for the faster intrinsic nucleotide dissociation properties of G13D. Such a hybrid could be used to evaluate how faster nucleotide dissociation would influence signaling through the comparison of this hybrid’s behavior with that of the G12D mutant.

The Ras network within the colorectal cancer context is assumed to be EGFR driven, and EGFR is assumed to activate Ras via increased activation of Ras GEFs like SOS1. We use a tenfold increase in V_max_ for GEF reactions to indicate EGFR activation, just as we have done previously to model receptor tyrosine kinase mediated Ras activation (Stites et al., 2007). To simulate an EGFR inhibition dose response, levels of GEF activity between the “high” (10× increase) case and the basal “low” (1×) level were considered and the resulting level of RasGTP determined via model simulation. We assume that the three Ras proteins, HRAS, NRAS, and KRAS, share similar biochemistry and can be modeled with the same set of biochemical properties; such an assumption is consistent with measurements of the three Ras proteins (Ahmadian et al., 1997; Lenzen et al., 1998). We assume that measurements that provide α for one Ras protein are good approximations for the same mutant to the other Ras proteins. We assume that more than one Ras gene is expressed in colorectal cancer cells. This is consistent with many data (Mageean et al., 2015; Omerovic et al., 2008). We here model Ras mutants as being heterozygous, such that for a KRAS mutant, one half of total KRAS will be mutant and one half of total KRAS will be wild-type. Here, we assume that 50% of total Ras is KRAS (and that 25% of total Ras is mutant). This assumption is consistent with mass spectrometric quantification of KRAS, NRAS, and HRAS levels (Mageean et al., 2015).

RasGTP and RasGTP-effector complex are considered as measures of Ras pathway activation. Model simulations are used to determine steady-state levels of RasGTP and RasGTP-effector. Simulations and analysis are performed in MATLAB (9.1.0.441655, MathWorks).

### Western Blot Analysis

Cell lysates were generated using radioimmunoprecipitation assay (RIPA) buffer (150 mM NaCl, 1% nonyl phenoxypolyethoxylethanol (NP-40), 0.5% sodium deoxycholate, 0.1% sodium dodecyl sulfate, 50 mM Tris of pH 8.0) containing protease inhibitor cocktail (Cell Signaling Technologies, Dancers MA) and incubated on ice for 1 hour. Total protein concentration was determined by Pierce-Protein assay (Thermo Fisher Scientific, Waltham MA). Protein samples (20 μg) were resolved by electrophoresis on 10-12% sodium dodecyl sulfate–polyacrylamide gels and electrophoretically transferred to polyvinylidene difluoride (PVDF) membranes (Millipore Corporation, Bedford, MA) for 20 minutes at 25V. The blots were probed with the appropriate primary antibody and the appropriate fluorophore conjugated secondary antibody. The protein bands were visualized using the Licor Clx Odyssey imaging station (Licor Biosystems, Lincoln NE). Comparative changes were measured with Licor Image Studio software.

### Cell-Proliferation Assay

Cells (5000 per well) were seeded in 96-well plates in phenol-red-free medium supplemented with charcoal-stripped FBS. Treatments were initiated after the cells were attached. At the appropriate time points, cell viability was determined by MTT assay; 10 μl of MTT (5mg/ml in phosphate-buffered saline) was added to each well followed by incubation at 37°C for 2 hours. The formazan crystal sediments were dissolved in 100 μl of dimethyl sulfoxide and absorbance was measured at 590 nm using the Tecan Infinite 200 Pro-plate reader (Tecan, Mannedorf, Switzerland). Each treatment was performed in seven replicate wells and repeated three times.

### Colony-Formation Assay

Cells were trypsinized and 4000 cells per well were plated in triplicate 6-well plates in DMEM supplemented with FBS. Colonies were formed after 7 days. The cells were fixed with ice-cold methanol and stained with crystal violet. Images were obtained using the Licor Clx Odyssey imaging station (Licor Biosystems, Lincoln NE). Colony formation was quantified by measuring absorbance per well. Comparison were made by normalizing to control wells. A total of five experimental replicates were performed with each containing three technical replicates.

### Active Ras Pulldown Assay

Isolation of active GTP bound Ras was performed using the Active Ras Pull-Down and Detection Kit (Thermo Fisher Scientific, Waltham MA) following manufacturers protocol. Ras abundance was measured by Western blot and/or by mass spectrometry.

### Mass Spectrometry

Samples were precipitated using Methanol-Chloroform. Dried pellets were dissolved in 8 M urea, reduced with 5 mM tris (2-carboxyethyl) phosphine hydrochloride (TCEP), and alkylated with 50 mM chloroacetamide. Proteins were then trypsin digested overnight at 37° C. Samples were digested at 50 μl final volume. Heavy labeled peptides were spiked-in to the digested samples at appropriate concentrations so that a single LCMS injection contained 10 μl of digested sample with 500 fmol of heavy labeled peptides. The samples were analyzed on a Fusion mass spectrometer (Thermo). Samples were injected directly onto a 25 cm, 100 μm ID column packed with BEH 1.7 μm C18 resin (Waters). Samples were separated at a flow rate of 300 nL/min on a nLC 1200 (Thermo). Buffer A and B were 0.1% formic acid in water and 90% acetonitrile, respectively. A gradient of 1–25% B over 110 min, an increase to 40% B over 10 min, an increase to 100% B over another 10 min and held at 90% B for a final 10 min of washing was used for 140 min total run time. Peptides were eluted directly from the tip of the column and nanosprayed directly into the mass spectrometer by application of 2.8 kV voltage at the back of the column. The Fusion was operated in a data dependent mode. Full MS1 scans were collected in the Orbitrap at 120k resolution. The cycle time was set to 3 s, and within this 3 s the most abundant ions per scan were selected for CID MS/MS in the ion trap. Monoisotopic precursor selection was enabled and dynamic exclusion was used with exclusion duration of 5 s

Peak area quantitation of the heavy peptides and corresponding light peptides from the samples were extracted by Skyline (MacLean et al., 2010). Within each sample, we used mutant Ras as a standard to normalize against. We then compared the ratio of normalized wild-type peptide levels in cetuximab treated conditions to normalized wild-type peptide levels in non-cetuximab treated conditions.

### siRNA-mediated Gene Knockdown

The appropriate recombinant SW48 cells were plated in a 10cm plate in DMEM supplemented with 10% FBS 24 h before transfection. The following day, cells were transfected with siRNAs against NF1 (2 μg) or control siRNA (2 μg) using Lipofectamine 2000.

### Expression Plasmid Transfection

Cells were plated in 96 well plate at 5000 cells per well in antibiotic free media. 24 h later cells were transfected with expression plasmids with duplex containing 0.2 μg of DNA and 0.25ul of Lipofectamine 2000 per well. Cell proliferation was assayed within at least 48 h.

### Co-immunoprecipitation

H293T cells were individually transfected with the expression plasmid for NF1-Flag, WT KRAS-GFP, G12V KRAS-GFP, G12D KRAS-GFP or KRAS G13D-GFP. Cells were harvested in IP Lysis/Wash Buffer (0.025M Tris, 0.15M NaCl, 0.001M EDTA, 1% NP-40, 5% glycerol; pH 7.4 and 1× protease inhibitor) 24 h post-transfection. Whole cell lysates (500 μg) were precleared for 0.5 h using Control Agarose Resin slurry (Thermo Fisher Scientific, Waltham MA). Immunoprecipitation was performed by first incubating 800 μl of H293T NF1-Flag pre-cleared lysate with 200 μl of either WT KRAS-GFP, G12V KRAS-GFP, G12D KRAS-GFP or G13D KRAS -GFP pre-cleared cell lysate. Each cell lysate mixture had EDTA (pH 8.0) added to make a final concentration of 10mM. GTP-gamma-S was added to the solution to a final concentration of 100nM. This solution was incubated at room temperature for 20 minutes with gentle rocking. The reaction was terminated by adding MgCl_2_ to the solution at a final concentration of 50mM. The final steps of the Co-IP were performed using the Pierce immunoprecipitation Kit (Thermo Fisher Scientific, Waltham MA) with immobilized anti-NF1 Ab (Santa Cruz Biotechnologies, CA). 500 μg of the cell lysate was added and incubated at room temperature under rotary agitation for 2 h. At the end of the incubation, the complexes were washed five times with Lysis buffer. The Western blotting was probed with mouse monoclonal NF1 antibody (Santa Cruz Biotechnologies, CA) and mouse monoclonal RAS antibody (Thermo Fisher Scientific, Waltham MA).

### Statistical Analysis

Significant differences amongst sample groups were determined by one-way ANOVA followed by post-hoc Tukey’s test for multiple comparisons with GraphPad Prism7 software. Mass spectrometry was performed twice. Every other experiment was performed at least three times, and P values are indicated in each figure.

## Acknowledgments

We thank Shumei Kato for providing cetuximab.

## Author contributions

T.M, J.M., A.S., and E.S. designed the experiments. T.M., M.M., J.D., S.L., K.P., and L.S-H. performed the experiments. T.M., J.D., and E.S. analyzed the experimental data. E.S. performed the computational analysis and wrote the manuscript with input from the other authors.

